# Cellular morphology emerges from polygenic, distributed transcriptional variation

**DOI:** 10.64898/2026.03.12.711281

**Authors:** Seyedezahra Paylakhi, Rafael Geurgas, Antionette Yasko, Robbee Wedow, Matthew Tegtmeyer

**Author notes:** Senior Author(s) (Robbee Wedow, Matthew Tegtmeyer).

## Abstract

Height and disease risk are canonical polygenic traits, but whether an analogous architecture governs cellular phenotypes remains unclear. Here, we show that cellular morphology behaves as a polygenic systems phenotype by integrating cross-modal modeling, perturbation profiling, and human genetic variation. An adversarially aligned autoencoder trained on four perturbation datasets predicted morphology from gene expression and generalized without retraining to matched RNA-seq and Cell Painting profiles from 100 genetically diverse iPSC donors. Seventeen morphological features were strongly predicted, particularly those capturing organelle distribution and cytoskeletal architecture. Predictive signal was broadly distributed across genes, with weak individual gene–morphology correlations. Independent CRISPR perturbations supported model-prioritized genes including *TIAM1*, *RAB31*, and *ABCC5*, while genetic analyses identified variant–gene–morphology associations involving *PDHX*, *SLC11A2*, and *MAPKAPK5*. These findings extend polygenic models of complex traits to cellular architecture and provide a framework for interpreting cross-modal prediction in functional genomics.

**Highlights:**

- Cellular morphology is encoded by broadly distributed transcriptional variation
- Transcriptome-to-morphology relationships generalize from perturbations to human donors
- CRISPR perturbations validate molecular anchors within distributed transcriptional programs
- Human genetic variation links regulatory genes to mitochondrial and cellular architecture

## Introduction

At the organismal level, complex traits, such as height, body mass index, and disease risk arise from the distributed influence of many loci rather than a handful of large-effect genes^1^. In the omnigenic model, even genes without direct mechanistic roles can influence phenotypic variation through interconnected regulatory networks. At the cellular level, transcriptional programs are translated into physical architecture through coordinated regulation of cytoskeletal organization, organelle positioning, and membrane dynamics. Whether this relationship is dominated by strong gene–phenotype effects or instead emerges from the integration of many weak transcriptional signals remains unresolved.

Cellular state can be measured through complementary molecular and morphological readouts. Transcriptomic profiles capture regulatory and pathway activity, whereas image-based assays such as Cell Painting quantify the integrated structural consequences of molecular, metabolic, and biophysical processes across cellular compartments^2–4^. These relationships are increasingly exploited in functional genomics and drug discovery, where compounds sharing mechanisms of action can produce similar morphological profiles^5,6^.

Large-scale multimodal datasets now measure gene expression and cellular morphology across thousands of perturbations^4,7–9^, enabling computational models to predict one modality from another with increasing accuracy. Approaches range from linear dimensionality reduction to nonlinear generative and foundation-model frameworks^10–14^. Yet high cross-modal accuracy leaves an important biological question unresolved: does prediction reflect a small number of interpretable gene–phenotype relationships, or the integration of many individually weak transcriptional signals?

Here, we test whether cellular morphology behaves as a polygenic trait by integrating cross-modal prediction with perturbational and human genetic analyses. Building on prior work characterizing morphological and transcriptional variation across human donors^15,16^, we train an adversarially aligned cross-modal model on four large perturbation datasets^7^ and test whether the learned transcriptional–morphological relationships generalize to natural inter-individual variation. We then determine how predictive signal is distributed across genes, test model-prioritized genes using independent CRISPR perturbation data, and connect these genes to cis-regulated expression through transcriptome-wide association analyses.

We find that highly predictable morphological features, particularly those describing organelle distribution and cytoskeletal architecture, arise from broadly distributed transcriptional information rather than strong single-gene associations. Independent CRISPR perturbations identify specific molecular anchors within these distributed programs, while genetic analyses provide complementary support for variant–gene–morphology relationships. Together, these results support cellular morphology as a polygenic systems phenotype and provide a framework for interpreting gene-level information from cross-modal models.

## Results

### Transcriptional programs encode large-scale cellular architecture across perturbations

To investigate how transcriptional variation shapes cellular structure, we analyzed multimodal perturbation datasets containing matched gene expression and Cell Painting morphological profiles. We applied a shared latent representation framework^11^ to four large-scale perturbation resources^7^: LUAD (lung adenocarcinoma–associated genetic perturbations), LINCS (Library of Integrated Network-Based Cellular Signatures pharmacological perturbations), TAORF (transient overexpression), and CDRP-bio (bioactive chemical perturbations). They quantify transcriptional state using the L1000 platform (∼978 landmark genes) alongside ∼1,000 CellProfiler^18^-derived morphology features per perturbation.

The modeling framework was adapted from the adversarial cross-domain translation architecture of Yang et al.^11^ (**Figure 1**). It consists of modality-specific linear autoencoders for gene expression and morphology, each projecting its input into a 150-dimensional latent representation. The morphology autoencoder was pretrained and subsequently frozen, while the RNA autoencoder was trained using RNA reconstruction and adversarial latent-distribution alignment. After alignment, RNA-derived latent representations were passed through the frozen morphology decoder to predict Cell Painting features.

**Figure 1.**
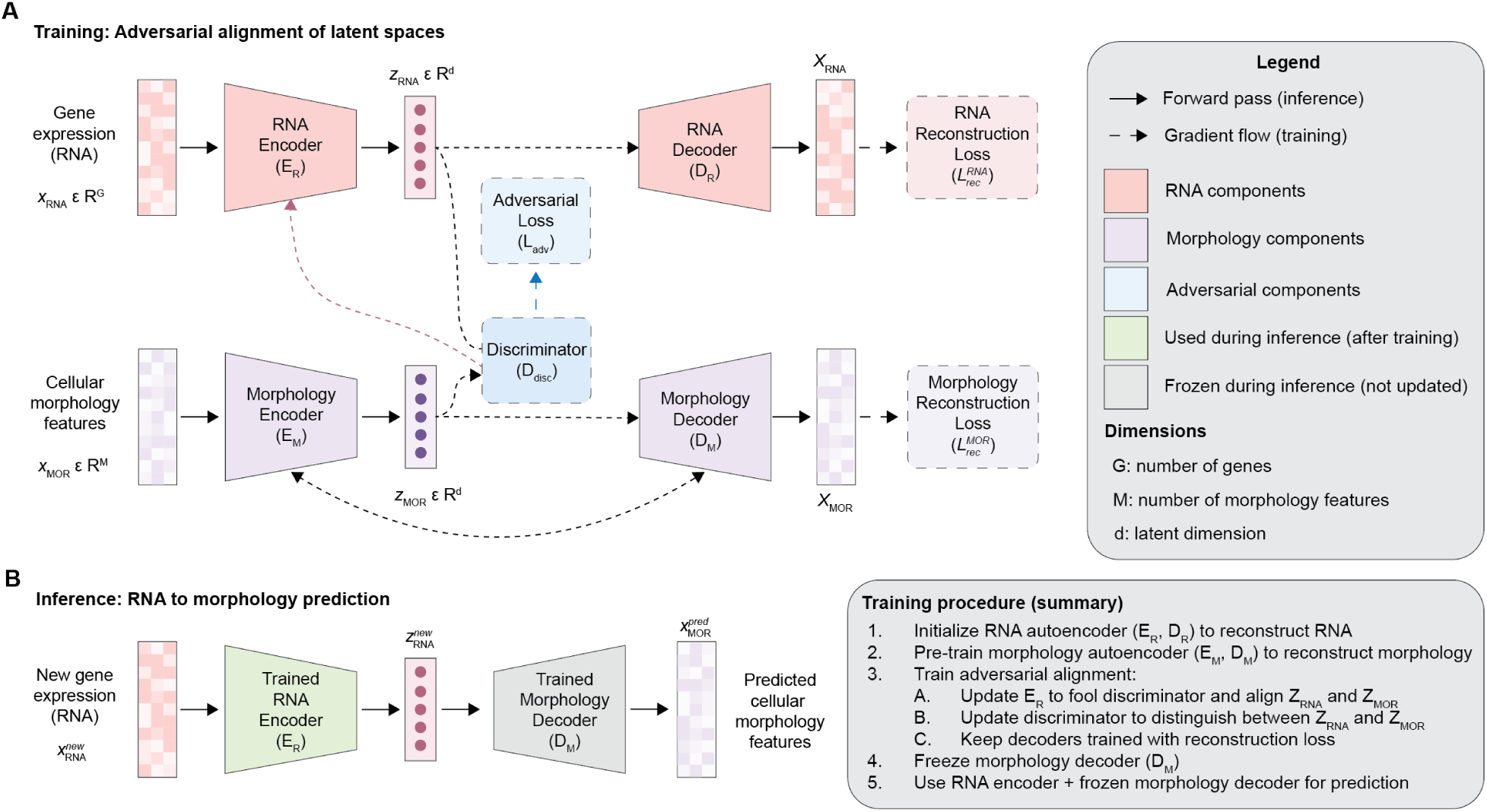
Schematic of the adversarially aligned cross-modal framework. The morphology autoencoder is pretrained and subsequently frozen. The RNA autoencoder is trained using an RNA reconstruction loss and an adversarial latent alignment objective. A discriminator aligns the distribution of RNA-derived latent representations with the pretrained morphology latent distribution. Morphology predictions are generated by decoding aligned RNA latent representations with the frozen morphology decoder.

Across all datasets, reconstruction loss decreased and stabilized during training (**Figure S1**), with final reconstruction errors indicating that both transcriptomic and morphological variation was retained in the learned representations (**Figure 2a**).

**Figure 2.**
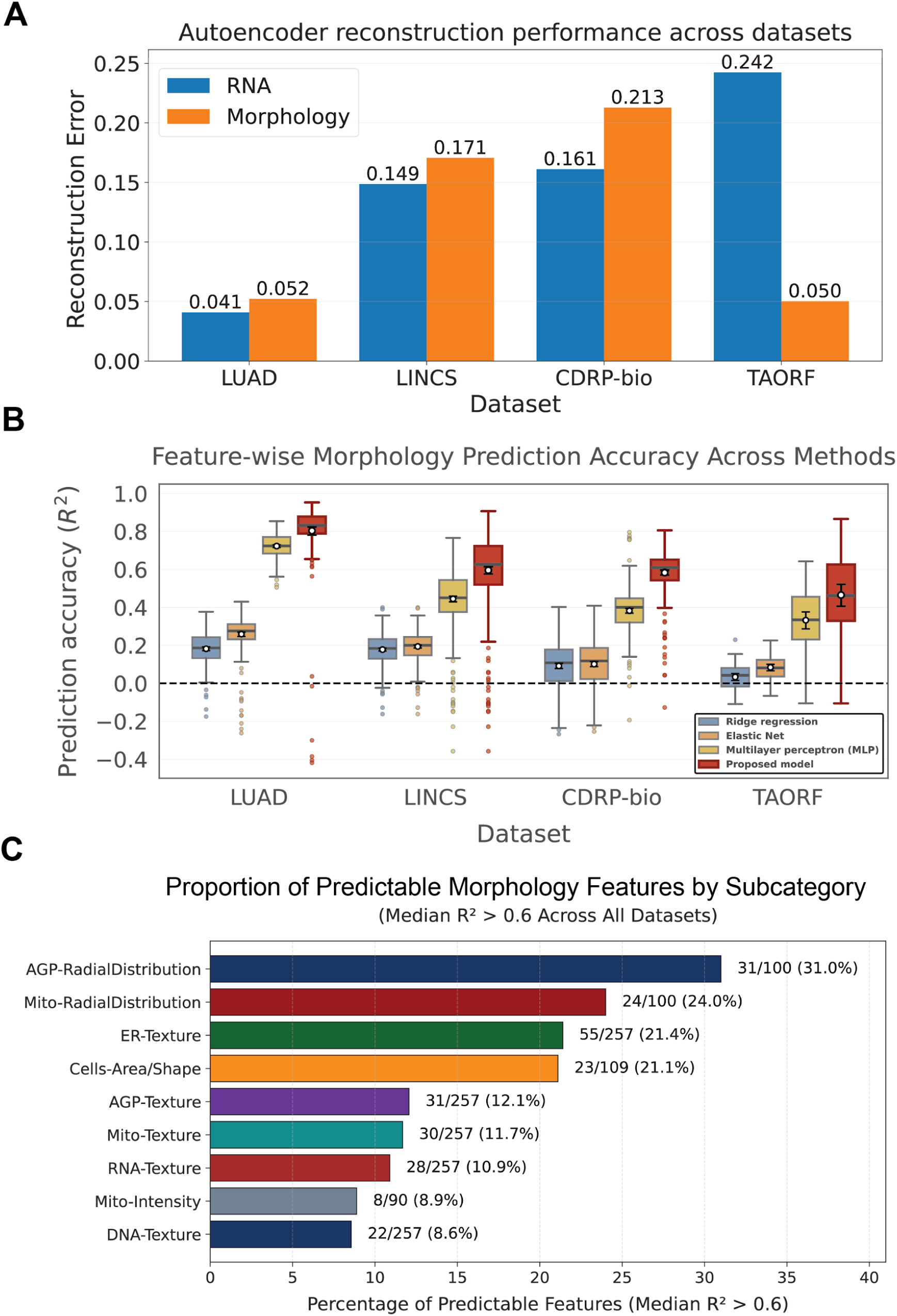
Performance of the proposed model across datasets and comparison with baseline methods. **(a)** Final held-out test mean squared reconstruction error (MSE) for the RNA and morphology autoencoders across four datasets (LUAD, LINCS, CDRP-bio, and TAORF). Bars represent modality-specific reconstruction performance on the held-out test set. (**b**) Comparison of feature-wise morphology prediction accuracy across four datasets using ridge regression, elastic net regression, a multilayer perceptron (MLP), and the proposed model. All methods were trained and evaluated using identical preprocessing, a fixed 70%/15%/15% train/validation/test split, and the same evaluation metric. Box plots show the distribution of prediction accuracy (R²) values across morphology features. White circles and error bars indicate the mean prediction accuracy and the corresponding 95% bootstrap confidence interval. (**c**) Bar plot showing the proportion of morphology features within each subcategory achieving a median R² > 0.6 across the four perturbation datasets. This pragmatic effect-size threshold was applied after model evaluation to define a focused subset of consistently predicted features for category-level enrichment and biological interpretation; it was not used during model training, hyperparameter selection, or model evaluation. Subcategories related to AGP radial distribution, mitochondrial radial distribution, and endoplasmic reticulum texture exhibited the strongest transcriptomic predictability.

We next evaluated whether transcriptional information encoded in the adversarially aligned latent representation could predict morphological features. All models were trained and evaluated using identical preprocessing, a fixed 70%/15%/15% train/validation/test split, and the same feature-level performance metric. Across all four datasets, the proposed model achieved robust prediction accuracy, with a substantial subset of morphology features exhibiting moderate to high predictability (**Figure 2b**). Compared with conventional machine-learning baselines, including ridge regression, elastic net regression, and a matched multilayer perceptron (MLP), the proposed model consistently achieved higher feature-level prediction accuracy.

Prediction accuracy collapsed when predicted profiles were randomly reassigned among held-out samples across 1,000 permutations (**Figure S2**), confirming that performance depended on the correct correspondence between transcriptomic and morphological profiles.

To focus downstream analyses on reproducibly predictable morphology, we selected features with a median R² > 0.6 across the four perturbation datasets. This reporting threshold was not used for model training or hyperparameter selection and identified features in the upper tail of the prediction distribution. For donor-level analyses, we additionally required permutation-based FDR q < 0.05.

These features were strongly enriched for AGP-RadialDistribution, Mito-RadialDistribution, and ER-Texture measurements (**Figure 2c**). In the Cell Painting assay, the AGP channel reports actin, Golgi, and plasma membrane organization, whereas the Mito channel captures mitochondrial distribution and abundance^3^. RadialDistribution features quantify how signal intensity is spatially distributed from the center of the cell toward the periphery, providing a quantitative readout of large-scale subcellular organization.

The enrichment of spatial organelle-distribution features suggests that transcriptional state is particularly coupled to global cellular organization. Rather than local descriptors such as nuclear shape or texture, the most predictable features captured the spatial arrangement of organelles and cytoskeletal structures across the cell.

### Transcriptional–morphological coupling generalizes to natural human genetic variation

Having established transcriptional prediction of morphology across experimental perturbations, we next tested whether these relationships extend to naturally occurring genetic variation. Because the LUAD perturbation data^7^ consistently achieved the highest predictive performance among the four perturbation datasets (**Figure 2b**), we selected the corresponding trained model to evaluate whether transcriptional–morphological relationships learned from a single perturbation dataset generalize to natural inter-individual variation. We then applied this model, without retraining or architectural modification, to predict morphology across 100 genetically diverse donors with matched RNA-seq and Cell Painting profiles on induced pluripotent stem cells (iPSCs) we previously described^15,17^. Donor transcriptomic profiles were restricted to the LUAD-trained L1000 gene set and processed using the preprocessing parameters learned from the LUAD training data.

This cross-context generalization, from pharmacological/genetic perturbation in A549 cells to natural inter-individual variation in iPSCs, represents a stringent test of whether transcriptional–morphological coupling reflects stable biological relationships or perturbation-specific statistical associations.

We evaluated transfer at two levels. Population-level performance measured, for each morphology feature, how well predicted values tracked inter-individual variation across the complete donor cohort. Individual-level prediction robustness was evaluated by nonparametric bootstrap resampling of individual donors. For each bootstrap resample, donors were sampled with replacement, feature-wise R² was recalculated using the same fixed model predictions, and the mean bootstrap R² was used as the individual-level prediction statistic for each morphology feature. Features with high population-level predictability generally also exhibited high individual-level prediction robustness (**Figure 3**). Because both the perturbation and donor datasets were represented using the L1000 landmark gene panel, which was specifically designed to capture broadly informative transcriptional variation, the observed cross-context generalization may in part reflect the informative nature of this curated feature set in addition to conserved biological relationships.

**Figure 3.**
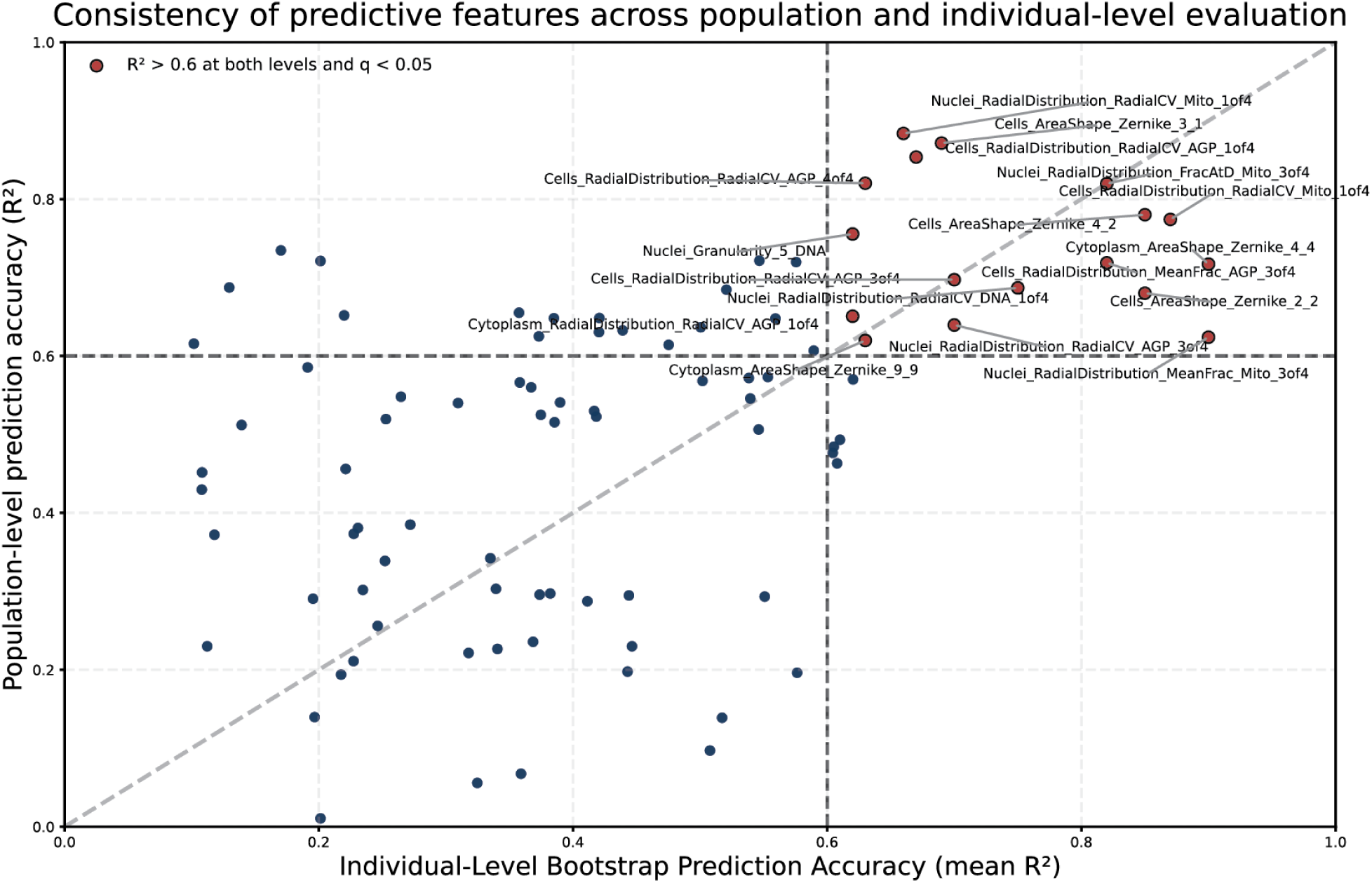
Comparison of global and individual-level morphology prediction Performance. Each point represents one morphology feature. The y-axis shows the global population-level prediction accuracy (feature-wise R² computed across the complete matched donor cohort; n = 100 donors). The x-axis shows the mean individual-level bootstrap prediction accuracy, calculated by repeatedly resampling individual donors with replacement and recomputing feature-wise R² using the same fixed model predictions. The dashed diagonal indicates equal global and bootstrap prediction accuracy. Features highlighted in red satisfied global R² > 0.6, mean individual-level bootstrap R² > 0.6, and permutation-based FDR q < 0.05.

Seventeen morphology features satisfied stringent effect-size and statistical significance thresholds (population-level R² > 0.6, mean individual-level bootstrap R² > 0.6, and permutation-based FDR q < 0.05; **Table S1**). Sensitivity analysis across R² thresholds of 0.5–0.8 retained 29, 17, 5, and 1 features, respectively (**Table S2**), supporting R² > 0.6 as a useful balance between predictive strength and feature retention. Ten of the 17 features were AGP- or Mito-RadialDistribution measurements. Thus, spatial organelle organization was among the most consistently predictable aspects of morphology across both perturbational and natural inter-individual variation.

### Predictable morphology reflects distributed transcriptional contributions

We next asked whether strong morphology prediction arises from a small number of dominant gene–phenotype relationships or from the distributed integration of many transcriptional signals. A sparse architecture should concentrate model importance in a small number of genes with strong marginal associations, whereas a polygenic architecture should produce distributed importance and weak individual gene–morphology relationships.

Gene contributions to each of the 17 highly predicted morphology features were quantified using effective end-to-end gene-to-morphology coefficients obtained by combining RNA encoder and morphology decoder weights.Model-derived importance scores were broadly distributed across genes, with no single gene dominating prediction of any of the 17 features (**Figure S3**). Consistent with this pattern, marginal gene–morphology correlations were uniformly weak (**Figure S4**).

This behavior is illustrated for two representative morphology features: Cells_RadialDistribution_RadialCV_AGP_1of4, which captures the variability of actin/Golgi/plasma membrane signal in the inner cytoplasmic quadrant, and Cells_AreaShape_Zernike_4_2, a Zernike-polynomial descriptor of cell geometry (**Figure 4**). For both features, predictive contributions were distributed across many genes while top-ranked genes displayed weak individual correlations (r < 0.2) with morphology. Thus, high multivariate importance did not require a strong univariate association.

**Figure 4.**
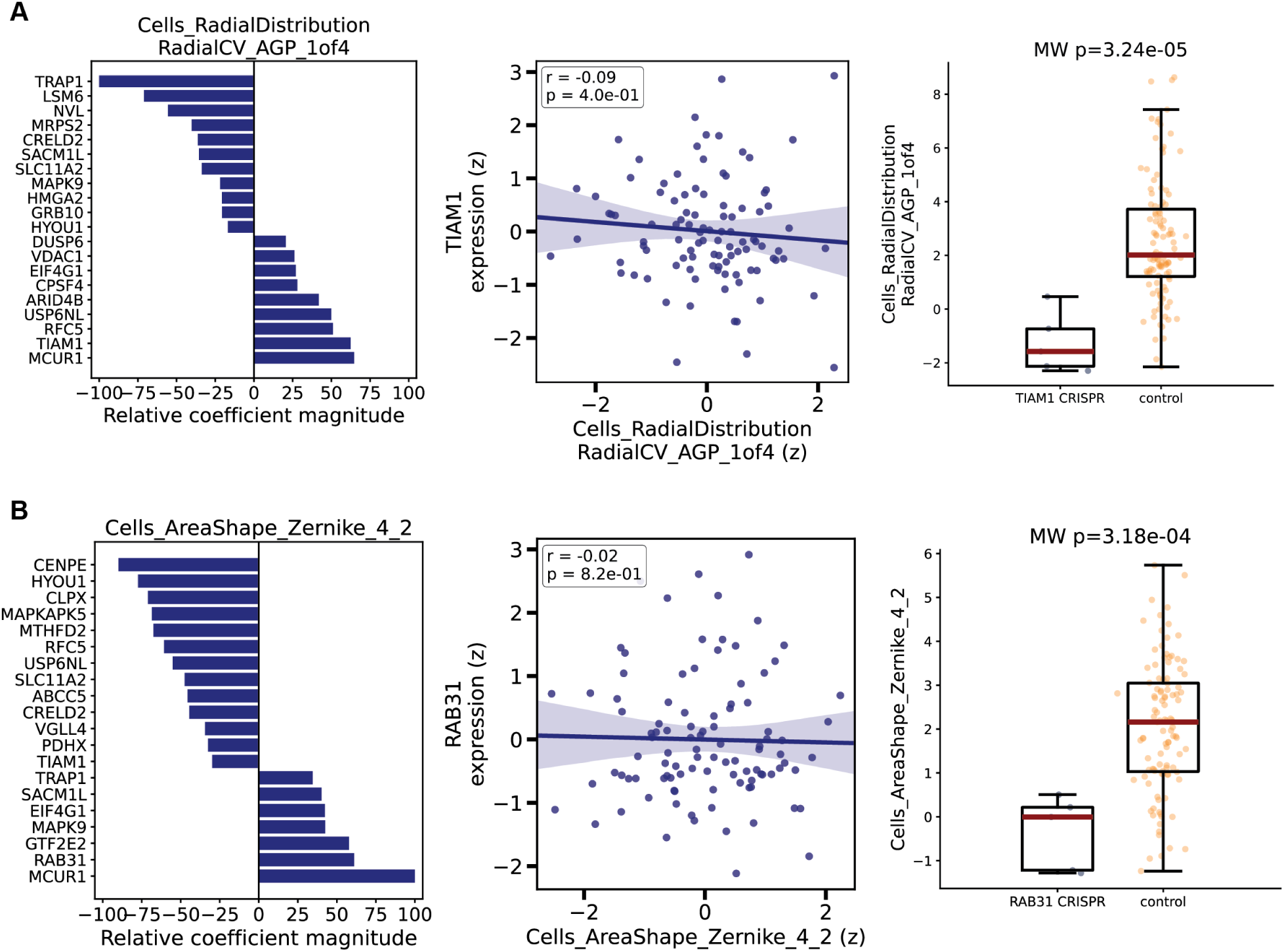
Distributed transcriptional prediction and CRISPR validation of representative gene–morphology relationships. **(A)** Prediction and validation of Cells_RadialDistribution_RadialCV_AGP_1of4. Left, top 20 genes ranked by relative coefficient magnitude in the transcriptome-to-morphology model. Effective end-to-end gene-to-morphology coefficients were scaled to the largest absolute coefficient for the feature; bar length indicates relative model contribution and sign indicates the direction of the learned relationship. Center, expression of the representative top-ranked gene TIAM1 shows little marginal correlation with the morphology feature despite its contribution to multigene prediction. The shaded region indicates the 95% confidence interval. Right, independent validation in JUMP Cell Painting data demonstrates that CRISPR perturbation of TIAM1 significantly alters AGP radial distribution relative to plate-matched non-targeting controls. **(B)** Corresponding analysis for Cells_AreaShape_Zernike_4_2. Left, top 20 genes ranked by relative coefficient magnitude. Center, expression of the representative top-ranked gene RAB31 shows little marginal correlation with the morphology feature. Right, independent CRISPR perturbation of RAB31 shifts the predicted AreaShape Zernike feature relative to plate-matched non-targeting controls.

To test whether this pattern depended on the cross-modal model, we performed an incremental-R² analysis using Ridge regression. Genes were ranked within each training fold, and models were fit using progressively larger predictor sets. Across features, approximately two-thirds to four-fifths of the gene panel was required to recover 90% of full-model held-out R² (**Figure S5**), indicating that predictive information was not concentrated in a small gene subset.

A nonlinear MLP coupled with permutation importance yielded the same qualitative result: gene rankings differed between models, but predictive contributions remained broadly distributed and marginal gene–morphology correlations weak (**Figure S6**). These analyses suggest that the distributed pattern is not solely attributable to coefficient dispersion in the linear model, although they do not establish independent causal effects for each contributing gene.

Together, these results support a distributed, polygenic transcriptional architecture underlying predictable cellular morphology.

### Perturbational and genetic analyses identify candidate molecular anchors of polygenic morphology

To determine whether model-prioritized genes correspond to biologically meaningful regulators of cellular architecture, we evaluated gene-level predictions using independent CRISPR perturbation data from the JUMP Cell Painting consortium^2^. This dataset provides high-content morphology profiles following targeted gene disruption and enables orthogonal validation of gene–morphology relationships independent of transcription-based prediction.

We use molecular anchor operationally to denote a gene that is both prioritized for a morphology feature by the cross-modal model and whose independent CRISPR perturbation significantly alters the same or a closely corresponding feature. The term does not imply an omnigenic “core gene” or a principal causal regulator, but rather an experimentally supported connection between a distributed transcriptional program and cellular phenotype.

Across morphology features with strong and reproducible predictability, disruption of several highly ranked genes produced significant and feature-specific morphology changes relative to plate-matched non-targeting controls.

*TIAM1* ranked among the top 20 predictors of AGP radial distribution despite weak marginal association with the phenotype, and its CRISPR disruption significantly altered AGP radial distribution (**Figure 4A**). *TIAM1* encodes a Rac1-specific guanine nucleotide exchange factor that regulates actin remodeling and membrane protrusion ^18,19^, providing a biologically coherent connection to AGP spatial organization.

*RAB31* showed the same pattern for Zernike-based AreaShape features: high model prioritization, weak marginal association, and significant morphological effects following CRISPR disruption (**Figure 4B**). Its established role in trans-Golgi and endosomal trafficking provides a plausible connection to membrane-dependent cell geometry ^20^.

*ABCC5* similarly ranked among the top predictors of the mitochondrial feature Nuclei_RadialDistribution_FracAtD_Mito_3of4 despite only modest marginal correlation (r = 0.24); its CRISPR disruption significantly shifted the same feature (Mann–Whitney p = 5.01 × 10⁻⁶). *ABCC5* encodes MRP5, recently implicated in mitochondrial-associated membrane function and mitochondrial structural integrity^21^. This convergence provides independent perturbational support for the model-predicted *ABCC5*–mitochondrial morphology relationship.

Together, these experiments show that genes prioritized within a distributed predictive architecture can nevertheless exert feature-specific morphological effects. *TIAM1*, *RAB31*, and *ABCC5* span distinct processes, cytoskeletal remodeling, membrane trafficking, and mitochondrial-associated membrane transport, illustrating how diverse molecular pathways can converge on cellular architecture without any single gene dominating prediction.

### Genetic association analyses identify statistically supported links among variants, gene expression, and morphology

We next asked whether naturally occurring cis-regulatory variation provided independent support for relationships between gene expression and morphology. For each of the 17 highly predicted features, we tested expression–morphology associations among genes with significant cis-eQTLs in the donor cohort ^16^, adjusting for nuisance covariates (**Methods**).

Despite the limited donor sample (N = 100), 71% of significant gene–feature associations involved model-prioritized genes, compared with approximately 20–30% expected from the fraction of genes prioritized (**Figure 5a; Table S3**). This enrichment suggests that model-based rankings capture signals not evident from marginal expression–morphology tests alone.

**Figure 5.**
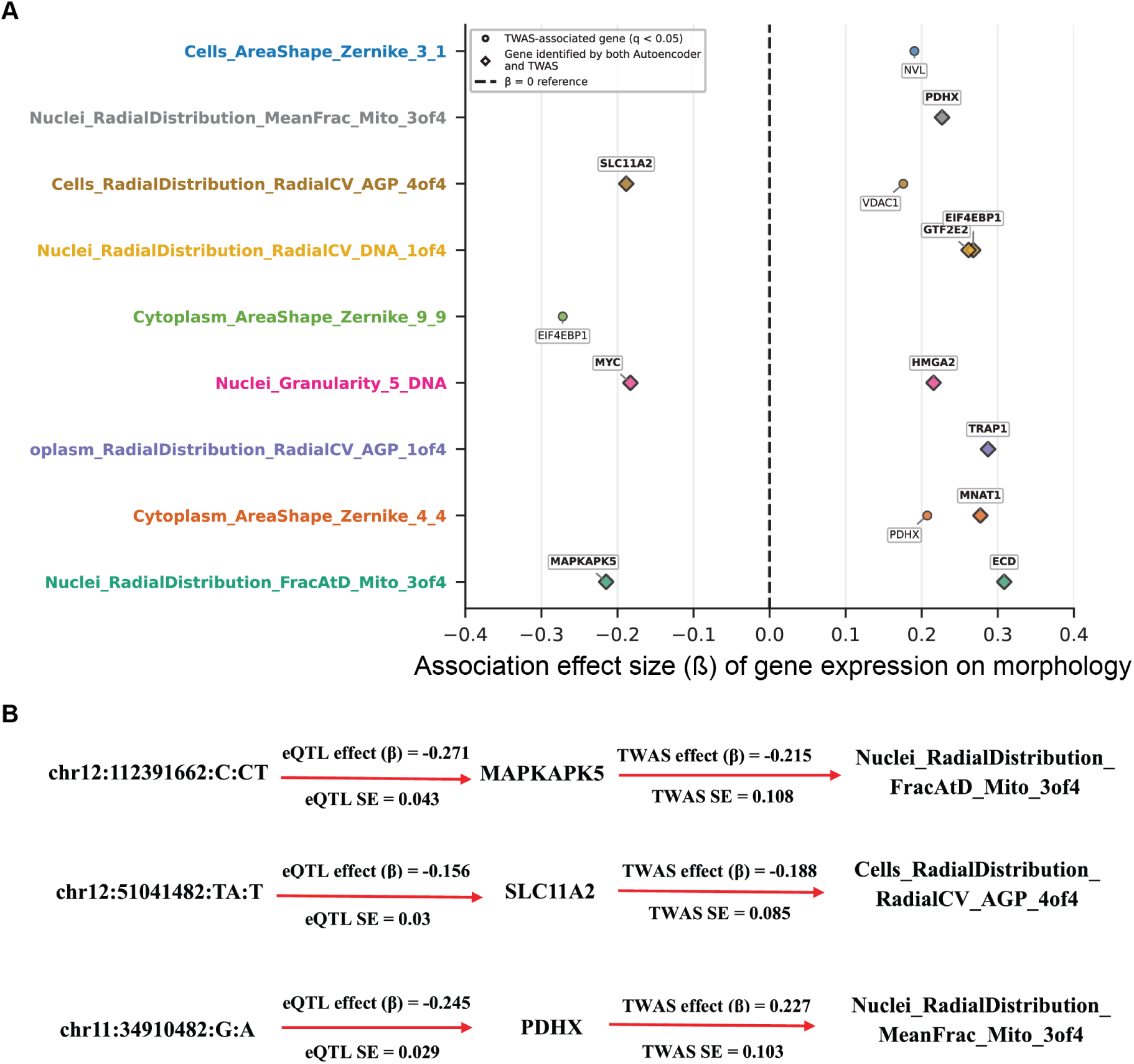
Convergence between predictive genes and genetically supported expression–morphology associations. (**A**) Overlap between genes prioritized by the shared autoencoder and genes identified by gene-level expression-morphology association analyses across top morphology features. Points show expression–morphology association effect sizes (β) for genes with cis-eQTL support.. Diamonds denote genes identified by both approaches. Associations are shown for genes passing false discovery rate correction (q < 0.05) within each morphology feature; the dashed line indicates β = 0. (**B**) Representative variant–gene–morphology chains linking genetic variation to morphology through gene expression. Each schematic depicts a significant cis-eQTL association (variant → gene expression; β ± SE) followed by a significant expression–morphology association (gene expression → morphology feature; β ± SE).. All illustrated eQTL and TWAS associations pass FDR correction (q < 0.05).

We then linked significant expression–morphology associations to each gene’s strongest cis-eQTL, yielding statistical variant–gene–morphology chains (**Figure 5b; Table S4**). Three representative associations illustrate biologically plausible links among genetic variation, gene expression, and cellular morphology:

PDHX encodes an essential component of the mitochondrial pyruvate dehydrogenase complex, which links glycolysis to the TCA cycle ^22^. Its association with Mito-RadialDistribution therefore identifies a plausible connection between genetically regulated mitochondrial metabolism and mitochondrial spatial organization, although the statistical chain does not establish mediation. SLC11A2 encodes the divalent metal transporter DMT1, a central regulator of cellular iron uptake and recycling ^23^. Its association with mitochondrial and membrane-related morphology is consistent with the dependence of mitochondrial metabolism on iron homeostasis. MAPKAPK5 encodes a stress-responsive kinase downstream of p38 MAPK that regulates cytoskeletal dynamics and cell motility ^24^, providing a plausible link to the spatial and cytoskeletal features enriched among highly predictable traits.

These associations provide limited but complementary genetic support for pathways spanning mitochondrial metabolism, iron homeostasis, and cytoskeletal regulation. Most model-prioritized genes lacked detectable genetic support, consistent with both the limited power of this cohort and a distributed architecture in which only a subset of contributing genes is expected to achieve statistical significance.

### Limitations of the study

First, the cross-modal model captures predictive associations rather than causal relationships. CRISPR experiments provide causal evidence for selected gene–morphology relationships, but the variant–gene–morphology chains do not establish mediation. Second, genetic analyses are limited by the donor sample size (N = 100) and the availability and context specificity of cis-eQTLs, reducing power to detect genetically regulated contributors. Third, model training and donor transfer were restricted to the ∼978 L1000 landmark genes. Because these genes were selected to capture broadly informative transcriptional variation, they may facilitate cross-context generalization, and it remains unclear whether the same distributed architecture would be observed using genome-wide expression. Fourth, we deliberately used a simple linear encoder–decoder architecture to favor interpretability over model complexity. More expressive architectures could potentially improve prediction, although the nonlinear MLP sensitivity analysis reproduced the same qualitative pattern of distributed gene importance and weak marginal associations. Fifth, batch correction was performed by the original publishers of each perturbation dataset and was not independently re-evaluated here; residual technical effects therefore remain possible. Finally, our analyses use well-level aggregate profiles and therefore cannot resolve cell-to-cell heterogeneity in transcriptional–morphological coupling. Matched single-cell transcriptomic and imaging measurements will be required to determine whether the degree of apparent polygenicity varies across cellular states or subpopulations.

## Discussion

This study addresses whether accurate prediction of cellular morphology from gene expression reflects direct gene–phenotype mappings or the integration of distributed transcriptional information. Across perturbational, genetic, and modeling analyses, we find that highly predictable morphological phenotypes are associated with many individually modest transcriptional contributions, while a subset of model-prioritized genes show feature-specific effects in independent CRISPR experiments. Together, these findings support a polygenic architecture for cellular morphology.

The central finding is that strong cross-modal prediction does not require strong univariate gene–morphology relationships. Across the 17 features that generalized to human donors, model importance was broadly distributed while individual gene–morphology correlations remained weak. Incremental-R² analyses further showed that most of the gene panel was required to recover full-model performance, and a nonlinear MLP reproduced the same qualitative pattern. Thus, the distributed architecture is unlikely to arise solely from coefficient dispersion in the linear model.

This organization parallels key features of the omnigenic model of organismal traits, in which regulatory networks distribute phenotypic influence across many genes rather than a small number of dominant loci ^25^. We do not infer that every model-prioritized gene independently causes morphology; instead, the results indicate that transcriptional information relevant to cellular architecture is distributed broadly across the expression state. Consequently, gene rankings from cross-modal models should not be interpreted as direct causal rankings without independent validation.

The transfer of a model trained in perturbed A549 cells to inter-individual variation in iPSCs further suggests that some transcriptional–morphological relationships are conserved across biological contexts. AGP- and Mito-RadialDistribution features were particularly reproducible, suggesting that global cytoskeletal and organelle organization may provide robust morphological readouts of transcriptional state^2,26^. This interpretation remains qualified by the use of the curated L1000 landmark panel, which may facilitate cross-context transfer.

CRISPR validation nevertheless demonstrates that specific genes embedded within these distributed programs can exert measurable, feature-specific morphological effects. *TIAM1*, *RAB31*, and *ABCC5* connect the model-derived predictions to cytoskeletal remodeling, membrane trafficking, and mitochondrial-associated membrane biology, respectively. Their mechanistic diversity reinforces a model in which multiple molecular processes converge on cellular architecture rather than a single pathway dominating morphology.

The variant–gene–morphology associations provide complementary evidence that a subset of these transcriptional relationships is coupled to inherited regulatory variation. *PDHX* and *SLC11A2*, for example, connect mitochondrial metabolism and iron homeostasis to morphology features involving mitochondrial or membrane organization. These chains do not establish causal mediation, but they generate experimentally testable hypotheses linking cis-regulatory variation to cellular architecture.

Most predictive genes lacked detectable genetic support, which is unsurprising given the modest donor sample (N = 100) and the contribution of environmental, stochastic, and trans-regulatory sources to gene-expression variation. Partial genetic anchoring is therefore compatible with a distributed architecture rather than evidence against it.

These findings have practical implications for interpreting cross-modal models. High predictive accuracy should not by itself be taken as evidence for direct mechanistic relationships between individual features across modalities: strong prediction can emerge from the integration of many weak and partially redundant molecular signals. Gene rankings should therefore be interpreted as candidate components of coordinated transcriptional programs and subjected to perturbational or genetic validation before causal conclusions are drawn.

More broadly, these findings provide a potential explanation for the effectiveness of morphology-based phenotypic screening. Because cellular morphology integrates cytoskeletal, organellar, metabolic, and membrane processes, weak distributed molecular perturbations may converge on detectable architectural phenotypes even when no individual molecular signal is dominant. A polygenic model of cellular morphology therefore provides a conceptual basis for the sensitivity of image-based profiling to diverse genetic and pharmacological perturbations ^5^.

Future work should determine whether this distributed architecture extends across cell types, molecular modalities, and genome-wide transcriptional measurements. Larger genetically diverse cohorts will also be required to distinguish direct from indirect variant–gene–morphology relationships. Extending these analyses to the single-cell level could further reveal whether transcriptional–morphological coupling varies across cellular subtypes or states. Specific cellular states may exhibit stronger genetic control of morphology or greater dependence on particular transcriptional programs, relationships that may be obscured by population-level measurements. Single-cell measurements could also distinguish whether distributed effects reflect many genes acting broadly across cells or more concentrated gene–phenotype relationships that emerge only within particular cellular states. Such analyses may therefore identify state-dependent genetic effects and clarify how cellular context shapes the penetrance of regulatory variation on morphology. More speculatively, treating morphology as a polygenic cellular phenotype raises the possibility of developing morphological polygenic scores that aggregate distributed molecular information into predictors of cellular state or function.

### Resource availability

#### Lead contact

Requests for further information and resources should be directed to and will be fulfilled by the lead contact, Matt Tegtmeyer.

#### Materials availability

This study did not generate new unique reagents

#### Data and code availability

All code to reproduce analyses and results is accessible at https://github.com/paylakhi/rna-morphology-autoencoder

## Acknowledgements

This work was supported by Purdue Office of Research Seed Funding, AnalytiXIN Fellowship funds to M.T. and R.W.

## Author Contributions

M.T., R.W. and S.P designed the study and wrote the manuscript with input from other authors. S.P led the analysis with assistance from M.T. R.G. and A.Y. provided input and edits to the manuscript. contributed to project management and sequencing. M.T. and R.W. supervised the study.

## Declaration of generative AI and AI-assisted technologies in the writing process

During the preparation of this work, the authors used ChatGPT (OpenAI) to assist with language editing. The authors reviewed and approved the final manuscript.

### Supplemental information titles and legends Figure titles and legends

## Supplementary Figures

**Supplementary Figure 1.**
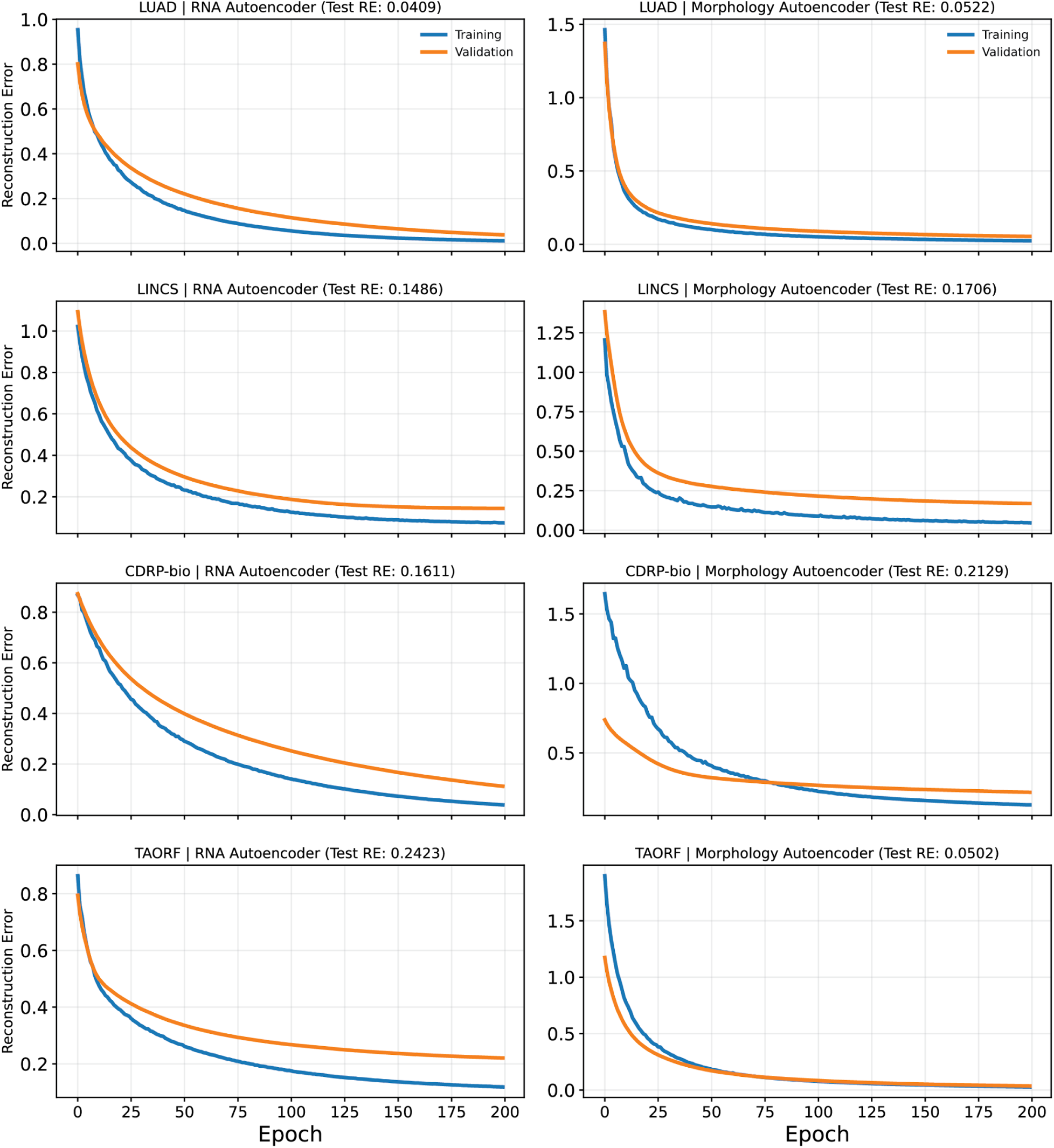
Training curves for RNA and morphology autoencoders. Training and validation reconstruction error across epochs for RNA (left) and morphology (right) autoencoders in LUAD, LINCS, CDRP-bio, and TAORF datasets. Reconstruction errors decrease and converge across epochs, indicating stable optimization. Final test reconstruction errors are shown in each panel.

**Supplementary Figure 2.**
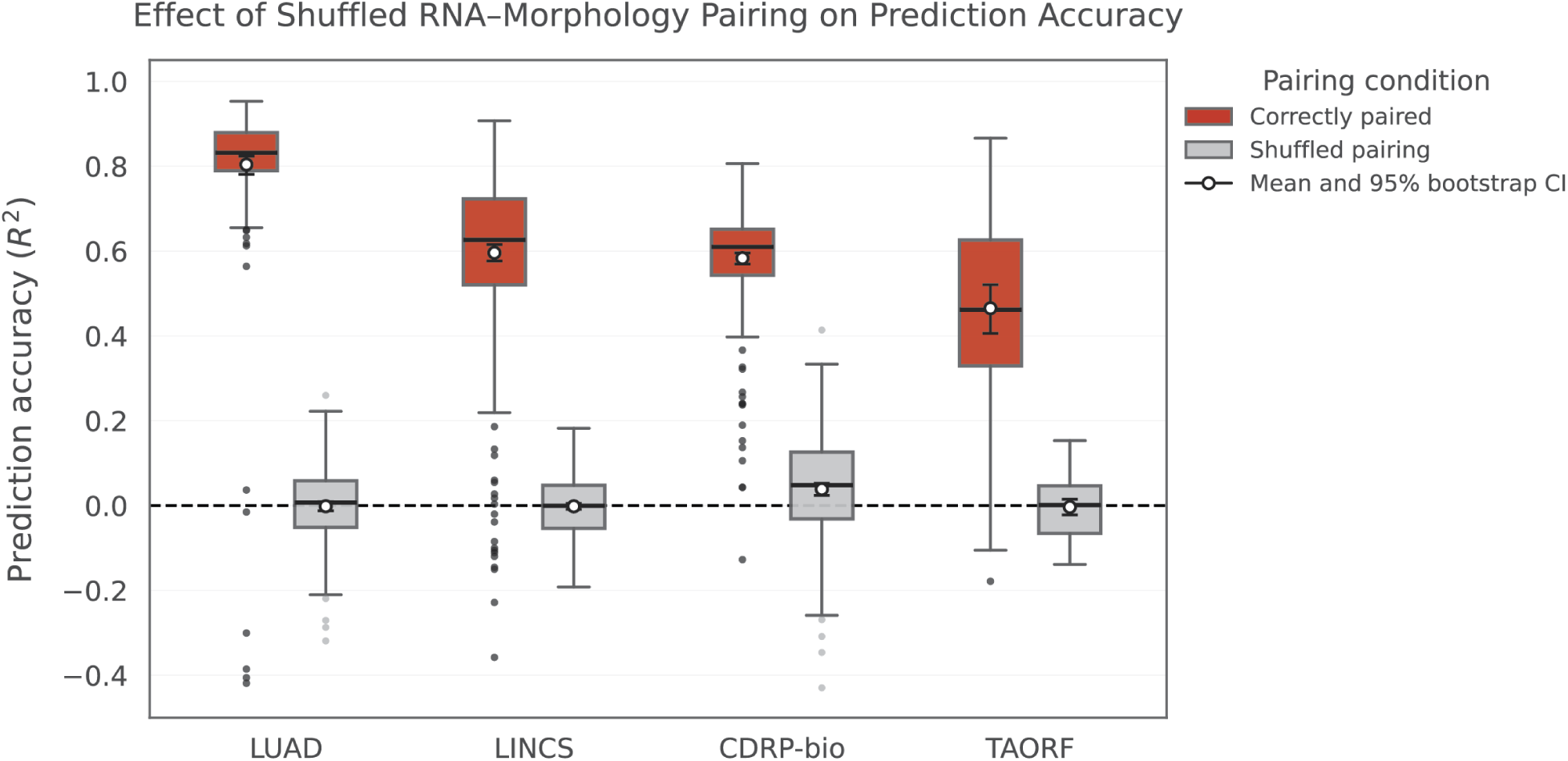
Effect of shuffled RNA–morphology pairing on morphology prediction accuracy. Box plots show the distributions of feature-level prediction accuracy obtained using the proposed model across the LUAD, LINCS, CDRP-bio, and TAORF datasets. Red boxes represent prediction performance evaluated using the original RNA–morphology correspondence in the held-out test set, whereas gray boxes represent a shuffled-pair control in which the previously generated predicted morphology profiles were randomly reassigned among held-out samples while the observed morphology profiles were kept fixed. This procedure preserves the distribution of predicted morphology while disrupting the correspondence between individual RNA and morphology samples. Feature-level R² values were recalculated across 1,000 random permutations. Center lines indicate medians, boxes indicate interquartile ranges, whiskers extend to 1.5× the interquartile range, and points beyond the whiskers indicate outlying morphology features. White circles and error bars indicate the mean prediction accuracy and corresponding 95% bootstrap confidence intervals. Shuffled pairing reduced prediction accuracy to approximately chance level across all four datasets, demonstrating that accurate morphology prediction depends on the correct RNA–morphology correspondence rather than spurious associations.

**Supplementary Figure 3.**
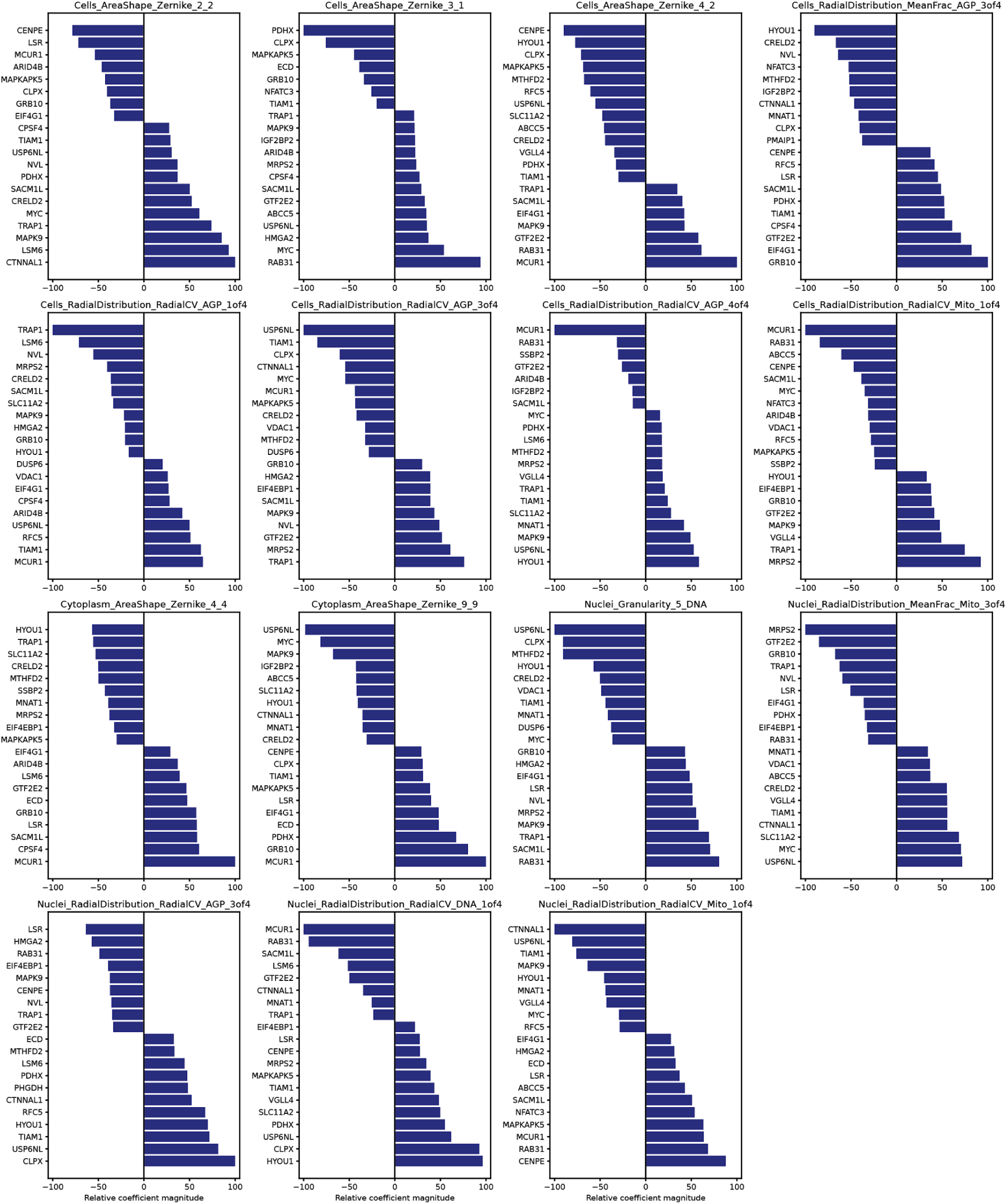
Distributed gene contributions underlying morphology prediction. For each representative morphology feature, the top-ranked genes by relative signed coefficient magnitude are shown. Both positive and negative contributions are observed across features and cellular compartments, highlighting that highly predictable morphology traits are driven by coordinated, multi-gene effects rather than strong univariate associations.

**Supplementary Figure 4.**
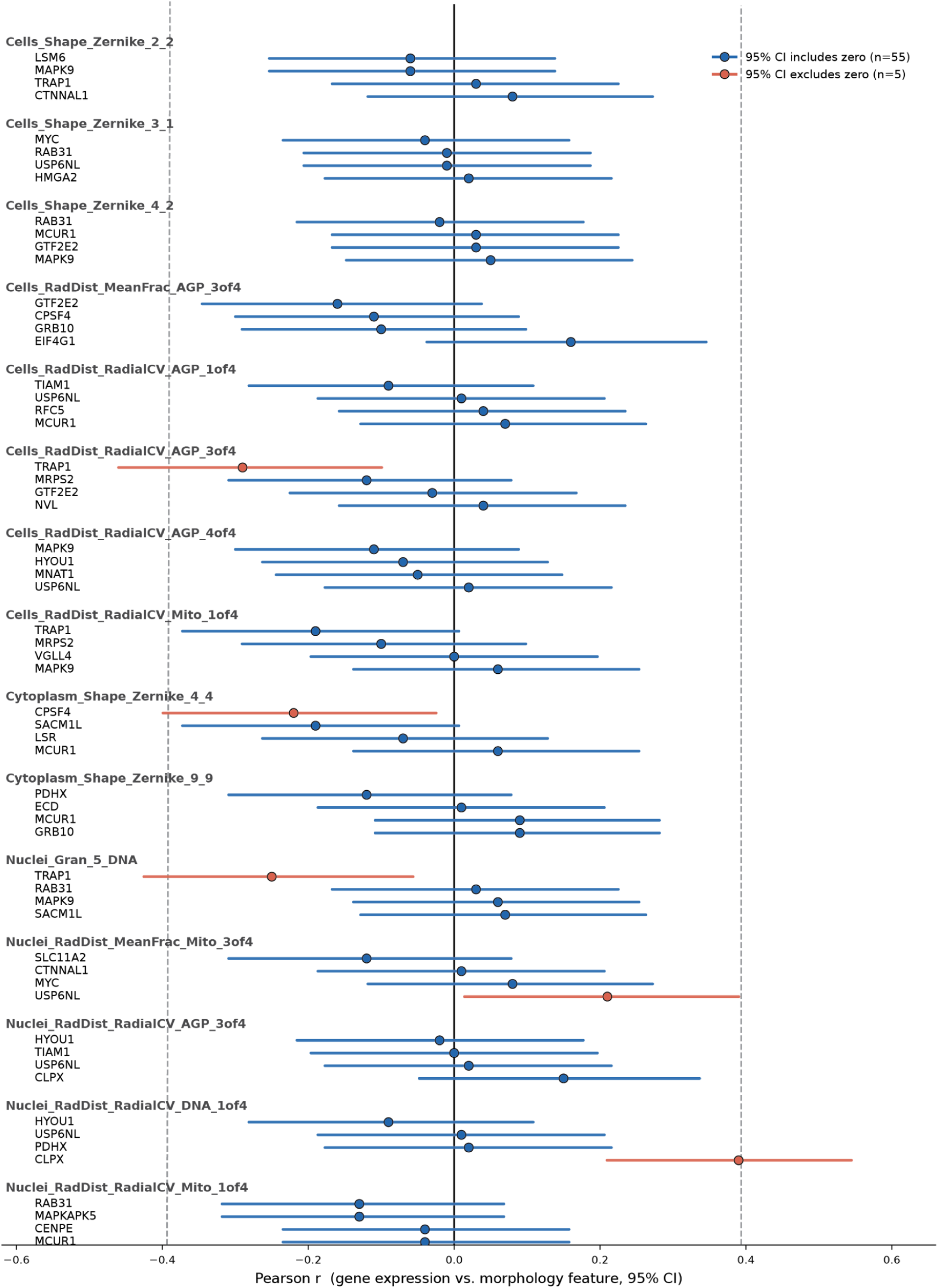
Marginal gene–morphology associations for model-identified genes. Forest plot shows Pearson correlations (*r*) between gene expression and morphology for model-prioritized gene–feature pairs. Points indicate Pearson *r* and horizontal lines indicate 95% confidence intervals. Pairs whose confidence intervals include zero are shown in blue, whereas those whose confidence intervals exclude zero are shown in red. Dashed vertical lines indicate the magnitude of correlation (|*r*| = 0.39) required to reach FDR *q* < 0.05. Across the 60 gene–morphology pairs, 55 had confidence intervals spanning zero and none reached the FDR significance threshold. The generally weak marginal associations, despite prioritization by the predictive model, support a distributed architecture in which morphology prediction arises from the integration of many modest transcriptional contributions rather than strong univariate gene–phenotype relationships.

**Supplementary Figure 5.**
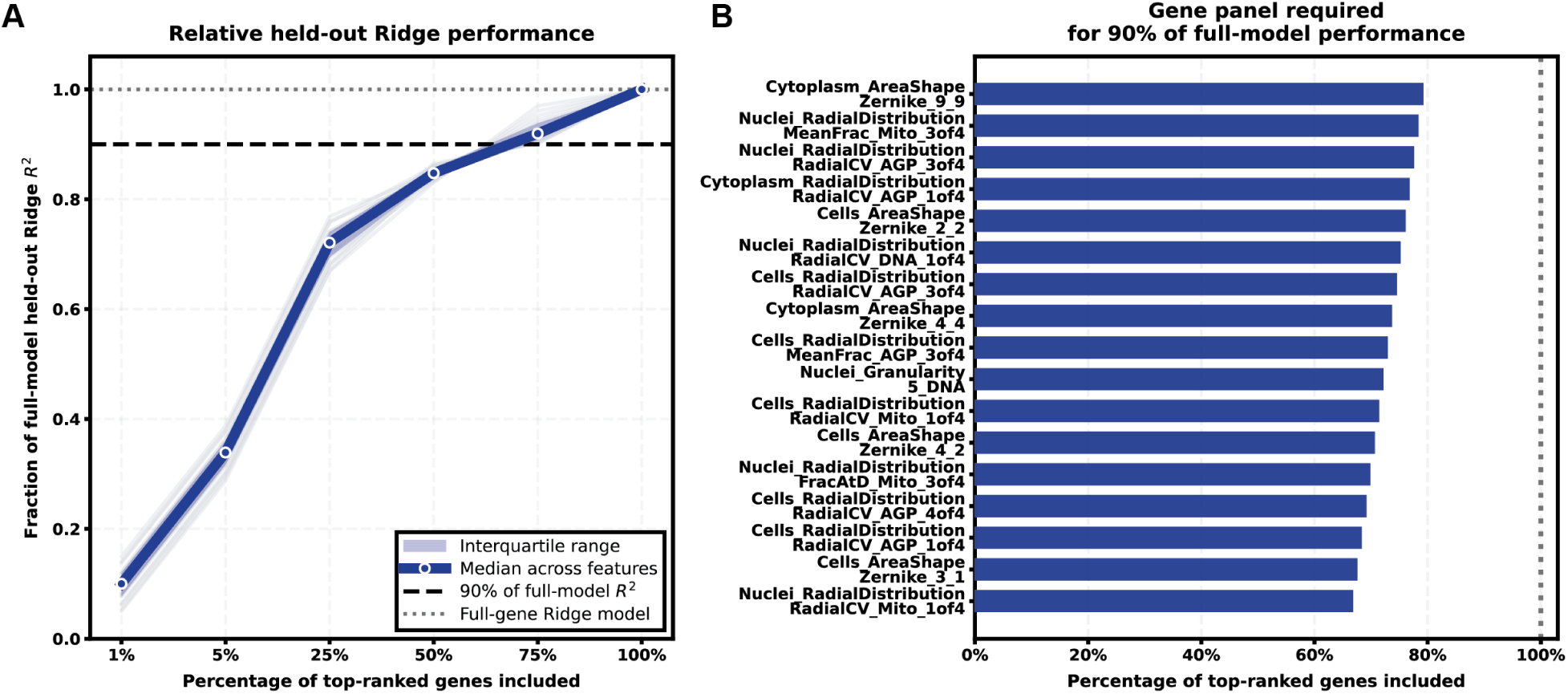
Incremental-R2 sparsity analysis of morphology prediction. (a) Relative held-out Ridge prediction performance as progressively larger fractions of the highest-ranked genes were included in the model. Within each cross-validation training fold, genes were ranked according to their association with the target morphology feature using only the training data. Ridge models were then fit using increasing fractions of the ranked gene panel and evaluated on held-out samples. The blue line represents the median relative performance across the 17 morphology features, the shaded region denotes the interquartile range, and faint gray lines show individual morphology features. The dashed horizontal line indicates 90% of the full-model held-out R2, while the dotted line indicates the full-gene model (100%). (b) Percentage of the ranked gene panel required for each morphology feature to recover 90% of the full-model held-out Ridge R2. Across features, approximately two-thirds to four-fifths of the ranked gene panel was required to approach full-model predictive performance, indicating that predictive information is distributed across many transcriptional features rather than concentrated in a small subset of dominant genes.

**Supplementary Figure 6.**
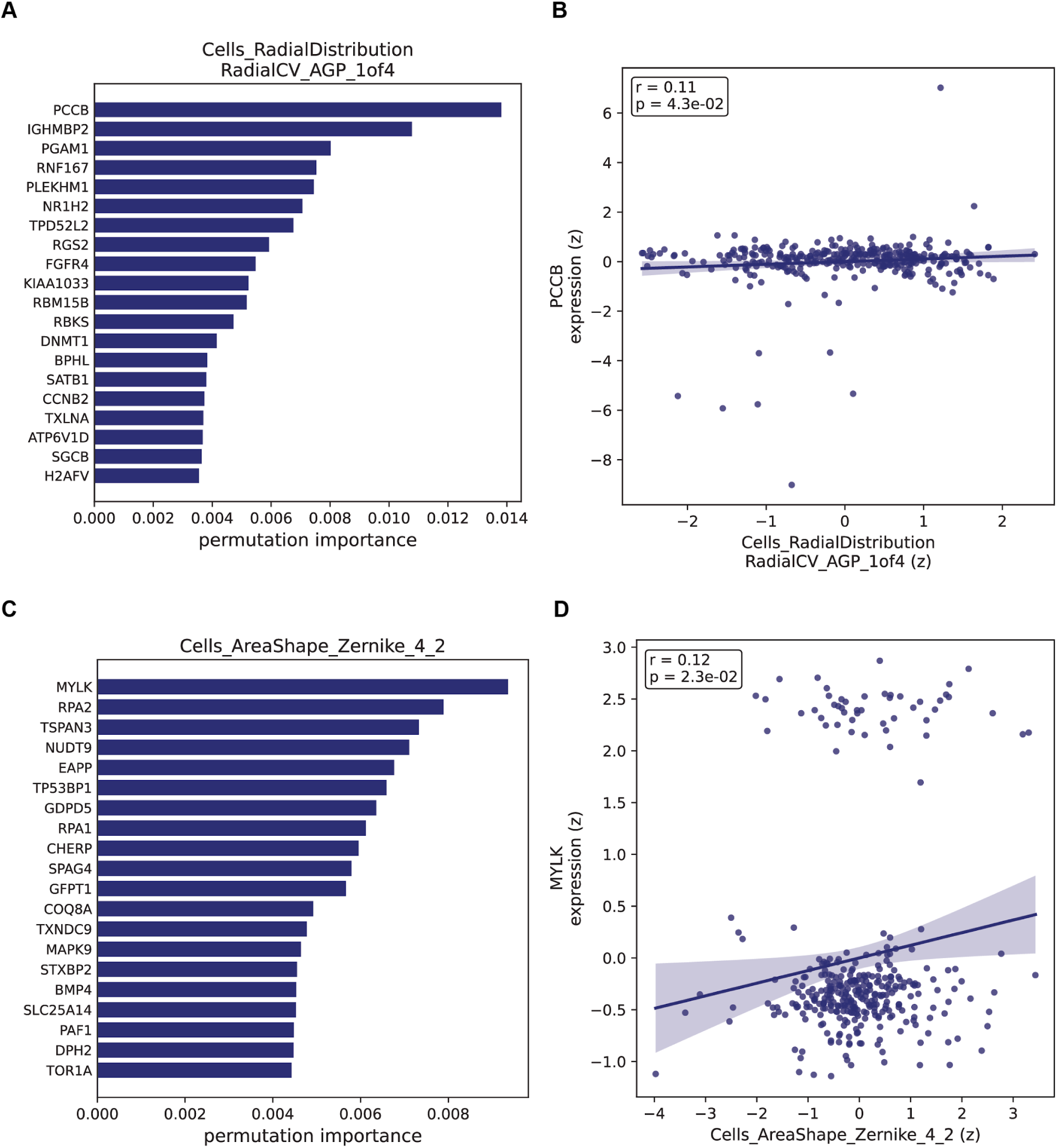
Nonlinear MLP sensitivity analysis of representative gene-morphology pairs. (a,c) Top 20 genes ranked by permutation importance for prediction of Cells_RadialDistribution_RadialCV_AGP_1of4 (a) and Cells_AreaShape_Zernicke_4_2 (c) using a nonlinear multilayer perceptron (MLP). Permutation importance indicates distributed predictive contributions across multiple genes rather than dominance by a single predictor. (b,d) Scatter plots of individual gene expression versus morphology for representative top-ranked genes, PCCB (b) and MYLK (d), demonstrate weak marginal correlations despite high multivariate predictive importance in the nonlinear MPL model. Shaded regions indicate 95% confidence intervals. Despite differences in model architecture and gene-importance estimation relative to the primary analysis, the nonlinear MLP similarly revealed distributed gene contributions and weak individual gene-morphology correlations, supporting the robustness of the observed polygenic architecture.

## Tables

**Supplementary Table 1. Highly predictable morphology features identified in donor-level transfer analysis.** Population-level prediction accuracy was quantified as the coefficient of determination (R²) for each morphology feature across the complete matched donor cohort. Ninety-five percent confidence intervals were estimated by bootstrap resampling. Robustness to donor composition was assessed using 5,000 nonparametric bootstrap resamples of donors, and the mean bootstrap R² is reported for each feature. Statistical significance was evaluated using 5,000 donor-label permutations followed by Benjamini–Hochberg false discovery rate correction. Morphology features shown here satisfied the predefined selection criteria of population-level R² > 0.6, mean bootstrap R² > 0.6, and FDR q < 0.05.

**Supplementary Table 2. Sensitivity of donor feature selection to the choice of effect-size threshold.** The final donor feature-selection procedure was repeated using joint population-level and mean bootstrap R² thresholds of 0.5, 0.6, 0.7, and 0.8 while retaining the permutation-based false discovery rate criterion (q < 0.05). For each threshold, the table reports the number of retained morphology features, the number of represented morphology categories, the median population-level and mean bootstrap R² values of the retained features, and the predominant morphology categories. The R² > 0.6 threshold provided a balance between predictive strength and sufficient feature representation for downstream biological interpretation.

**Supplementary Table 3. Significant expression–morphology associations for highly predictable morphology features.** Genes with significant cis-eQTL support were tested for association between gene expression and morphology across the donor cohort. The table reports morphology feature, gene, TWAS effect size (β), nominal p value, false discovery rate-adjusted q value, and whether the gene was prioritized by the cross-modal prediction model. Only associations passing Benjamini–Hochberg false discovery rate correction (q < 0.05) are shown.

**Supplementary Table 4. Statistically supported variant–gene–morphology associations.** Significant expression–morphology associations were linked to the strongest cis-eQTL for each gene to identify statistically supported variant–gene–morphology associations. The table reports the associated morphology feature, gene, lead cis-eQTL variant, eQTL effect size (β), standard error, p value and q value, together with the corresponding expression–morphology (TWAS) effect size, standard error, p value, and q value. These statistical associations provide complementary support for relationships among genetic variation, gene expression, and cellular morphology but should not be interpreted as evidence of causal mediation.

## STAR★METHODS

### RESOURCE AVAILABILITY

#### Lead contact

Further information and requests for resources should be directed to and will be fulfilled by the Lead Contact, **Matthew Tegtmeyer**.

#### Materials availability

This study did not generate new biological materials.

#### Data and code availability

This study analyzed previously published datasets together with computational analyses developed for this work.

1. Perturbation datasets are publicly available through their original repositories (see Key Resources Table).
2. Donor RNA-seq, Cell Painting, and cis-eQTL datasets are available as described in Tegtmeyer et al. and Wells et al.
3. Source code used for preprocessing, model training, statistical analyses, and figure generation is available at https://github.com/paylakhi/rna-morphology-autoencoder.
4. Any additional information required to reproduce this work is available from the Lead Contact upon request.

### EXPERIMENTAL MODEL AND STUDY PARTICIPANT DETAILS

#### Previously published perturbation datasets

This study analyzed four previously published multimodal perturbation datasets generated by Haghighi et al., including LUAD, LINCS, TAORF, and CDRP-bio. These datasets contain matched L1000 transcriptomic profiles and Cell Painting morphology measurements across pharmacological and genetic perturbations. No new perturbation experiments were performed. The datasets were obtained from the publicly available multimodal resource described by Haghighi et al.<u>(Haghighi et al. 2022)</u>. Preprocessed matched L1000 transcriptomic and Cell Painting morphology profiles are publicly available through the Broad Institute Rosetta resource.

### METHOD DETAILS

#### Data sources

We analyzed three classes of publicly available datasets: (i) previously published multimodal perturbation datasets containing matched L1000 transcriptomic and Cell Painting morphology profiles (LUAD, LINCS, TAORF, and CDRP-bio); (ii) previously published human donor-derived datasets containing matched RNA-seq, Cell Painting morphology, and cis-eQTL data from a subset of 100 human induced pluripotent stem cell (iPSC) donors; and (iii) independent CRISPR Cell Painting profiles from the JUMP Cell Painting Consortium (assembled CRISPR release v1.0a) for orthogonal validation of model-prioritized gene–morphology associations.

#### Data preprocessing

Sample identifiers were standardized within each dataset and matched across transcriptomic and morphological modalities using shared well identifiers. Duplicate well identifiers were removed prior to matching to retain a single paired transcriptomic and morphology profile for each shared well.

Matched samples were sorted to enforce identical row ordering across modalities before downstream analyses.

Only numeric features were analyzed. Infinite values were treated as missing. Features with ≥20% missing values were excluded, and remaining samples containing missing values were removed by complete-case filtering.

The final matched datasets contained 386 paired samples for LUAD, 360 for LINCS, 359 for TAORF, and 343 for CDRP-bio.

Raw RNA-seq count matrices were normalized using counts per million (CPM) followed by log1p transformation. Gene-expression filtering, scaling, and variance selection were estimated using the training subset only. Genes with mean training-set logCPM ≤ 5 were excluded, a StandardScaler was fitted on the remaining training-set gene-expression features, and genes with variance < 0.01 in the standardized training data were removed. The fitted scaler and variance selector were then applied unchanged to the validation and test subsets. Morphology correlation filtering and scaling were likewise estimated using the training subset only. Morphology features with absolute pairwise Pearson correlation > 0.90 in the training data were excluded, and a StandardScaler fitted on the remaining training morphology features was subsequently applied unchanged to the validation and test subsets.

#### Cross-modal model

The cross-modal framework was adapted from Yang et al. using modality-specific autoencoders for transcriptomic and morphological data.

Unlike the original nonlinear implementation, both encoder and decoder networks consisted of single fully connected linear layers to improve interpretability while preserving adversarial latent alignment.

Gene-expression and morphology data were projected into a shared 150-dimensional latent representation.

The morphology autoencoder was pretrained independently. Following pretraining, both the morphology encoder and decoder were frozen. During RNA training, latent representations produced by the RNA encoder were aligned with morphology latent representations through adversarial learning using a nonlinear discriminator.

The model was implemented in PyTorch.

The latent discriminator consisted of a fully connected layer mapping the 150-dimensional latent representation to 128 hidden units, followed by a LeakyReLU activation (negative slope = 0.2), dropout (p = 0.10), and a final fully connected layer producing a single discriminator logit.

The RNA and morphology encoders and decoders contained no nonlinear activation functions and each consisted of a single fully connected linear layer.

All linear layers were initialized using Xavier uniform initialization, with bias terms initialized to zero.

#### Model training

Training proceeded in two stages.

First, the morphology autoencoder was optimized using Adam (learning rate 1×10⁻⁴; weight decay 1×10⁻⁵) by minimizing morphology reconstruction loss.

Following convergence, morphology encoder and decoder parameters were frozen.

Second, the RNA autoencoder was trained using a weighted combination of RNA reconstruction loss and adversarial latent-distribution alignment loss:

*L_RNA_* = α*L_RNA_*_,*reconstruction*_ + λ*L_adv_* where RNA reconstruction loss was computed using mean squared error, adversarial loss encouraged RNA-derived latent representations to match the morphology latent distribution, and α = 0.1 and λ = 1.0.

The discriminator was optimized using binary cross-entropy loss:

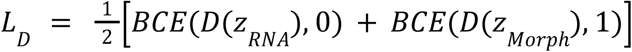

Both training stages used Adam (learning rate 1×10−4; weight decay 1×10−5) with a batch size of 16 for up to 200 epochs. Early stopping was based on validation loss, with a patience of 30 epochs during morphology-autoencoder pretraining and 40 epochs during RNA-adversarial alignment. The checkpoint with the lowest validation objective was retained for each training stage. All training used a fixed random seed of 42. One discriminator update was performed per RNA-autoencoder update, and gradient norms were clipped to 2.0.

#### Perturbation-level prediction

For each perturbation dataset, matched samples were randomly partitioned into training (70%), validation (15%), and test (15%) subsets using a fixed random seed. The validation subset was used for early stopping and model checkpoint selection, whereas the held-out test subset was used only for final evaluation. Morphology prediction was performed by encoding held-out RNA profiles with the trained RNA encoder and reconstructing morphology using the frozen pretrained morphology decoder. Prediction accuracy was quantified independently for each morphology feature using the coefficient of determination (R²). Morphology features with a median R² > 0.6 across the four perturbation datasets were designated highly predictable for downstream biological analyses.

#### Cross-context generalization

To evaluate generalization beyond perturbation experiments, the LUAD-trained model was applied directly to donor RNA-seq profiles without retraining or fine-tuning. The LUAD model was selected because it consistently achieved the highest prediction performance among the perturbation datasets (**Figure 2b**). Donor RNA-seq profiles were processed using the same preprocessing pipeline as the LUAD training data and restricted to genes overlapping the L1000 landmark panel. Donor morphology profiles were restricted to morphology features shared between the perturbation and donor datasets and evaluated in the same standardized morphology feature space as the LUAD-trained model. No donor-specific preprocessing parameters or model parameters were re-estimated.

Prediction performance was evaluated at two complementary levels. Population-level prediction accuracy was computed as the feature-wise coefficient of determination (R²) across the complete donor cohort. Individual-level prediction robustness was evaluated using 5,000 nonparametric bootstrap resamples of donors. For each bootstrap resample, donors were sampled with replacement, feature-wise R² was recalculated using the same fixed predicted and observed morphology profiles, and the mean bootstrap R² was reported for each morphology feature.

Population-level statistical significance was assessed independently using 5,000 donor-label permutations followed by Benjamini–Hochberg false discovery rate correction. Morphology features satisfying population-level R² > 0.6, mean bootstrap R² > 0.6, and permutation-based FDR q < 0.05 were considered consistently predictable.

To evaluate the sensitivity of feature selection to the choice of effect-size threshold, the donor-cohort feature-selection procedure was repeated using joint population-level and mean bootstrap R² thresholds of 0.5, 0.6, 0.7, and 0.8 while retaining the permutation-based FDR criterion (q < 0.05). For each threshold, we recorded the number of retained morphology features, the number of represented morphology categories, the median population-level and mean bootstrap R² values of the retained features, and the dominant morphology categories (**Table S2**).

#### Baseline prediction models

The adversarial autoencoder was compared with Ridge regression, Elastic Net regression, and a multilayer perceptron (MLP) using identical fixed training (70%), validation (15%), and test (15%) partitions and the same random seed (42). Hyperparameters were selected exclusively using validation performance, defined as the mean feature-level coefficient of determination (R²) across morphology features, and final models were retrained on the training data using the selected hyperparameters before evaluation on the held-out test set.

Ridge regression evaluated α values of {0.001, 0.01, 0.1, 1, 10, 100, 1000, 10000, 100000, and 1000000}, selecting the value that maximized validation performance.

Elastic Net used the MultiTaskElasticNet implementation with α values of {0.01, 0.1, 1, 10, 100} and l1_ratio values of {0.1, 0.5, 0.9}. The best parameter combination was selected using validation performance. Maximum iterations were set to 100,000 with a convergence tolerance of 1×10⁻⁴.

The nonlinear baseline used a single-hidden-layer multilayer perceptron consisting of 32 hidden units with ReLU activation and the Adam optimizer. Models were trained with batch size 8, learning rate 5×10⁻⁴, early stopping enabled (validation fraction = 0.20; n_iter_no_change = 50), and a maximum of 3000 iterations. L2 regularization strengths (α) of {0.001, 0.01, 0.1, and 1} were evaluated using the validation set.

#### Shuffled-pair control

To determine whether prediction accuracy reflected genuine transcriptome–morphology relationships rather than chance correspondence, predicted morphology profiles were randomly reassigned among held-out samples while observed morphology remained fixed.

Prediction accuracy was recalculated across 1,000 permutations.

#### Gene importance

For each morphology feature, gene importance was quantified using the effective end-to-end coefficient matrix of the linear RNA-to-morphology prediction pathway, calculated as ***W**_morphdecoder_ **W**_RNAencoder_*.

Effective coefficients were normalized within each morphology feature relative to the coefficient with the largest absolute magnitude.

Genes were ranked according to the absolute effective coefficient magnitude, whereas signed normalized coefficients were retained for visualization to distinguish positive and negative learned relationships.

#### Marginal correlations

For each morphology feature, Pearson correlation coefficients were computed between individual gene-expression values and morphology measurements across samples.

#### Incremental-R² analysis

To evaluate whether predictive information was concentrated in a small number of genes, genes were ranked independently within each outer training fold according to the absolute Pearson correlation between gene expression and each morphology feature.

Gene ranking was performed using training samples only to avoid information leakage into held-out test data.

Ridge regression models were then trained using progressively larger subsets of the ranked genes (1, 2, 5, 10, 20, 50, 100, 150, and all available genes).

For presentation, these subset sizes were expressed as approximate percentages of the complete gene panel, and relative held-out prediction performance was summarized as a function of the percentage of top-ranked genes included.

#### Nonlinear sensitivity analysis

To evaluate whether distributed gene importance depended on the linear model architecture, morphology prediction was repeated using a nonlinear multilayer perceptron (MLP).

For each of the 17 highly predictable morphology features, a separate MLPRegressor consisting of a single hidden layer with 32 hidden units, ReLU activation, Adam optimization, learning rate 5×10⁻⁴, batch size 8, early stopping, and L2 regularization (α = 0.1) was trained using five-fold cross-validation.

Gene importance was quantified using permutation importance computed on held-out test folds using 20 permutation repeats.

Mean permutation importance across cross-validation folds was used to rank genes for each morphology feature, and the highest-ranked genes were subsequently evaluated using marginal Pearson and Spearman correlations with morphology.

#### CRISPR perturbation validation

Independent CRISPR Cell Painting profiles from the JUMP consortium were used to provide orthogonal experimental validation of model-prioritized gene–morphology associations.

Morphology features corresponding to highly predicted features were matched between the perturbation and JUMP datasets based on identical Cell Painting feature names.

Representative genes (TIAM1, RAB31, and ABCC5) were selected because they ranked highly according to the model-derived gene importance scores for morphology features that were highly predictable by the cross-modal model.

For each predefined gene–feature pair, well-level CRISPR morphology profiles were merged with perturbation metadata using unique perturbation identifiers.

Negative-control wells were restricted to plate-matched non-targeting or no-guide controls originating from the same experimental plate(s) as the corresponding CRISPR perturbation.

Morphology measurements were compared between CRISPR-targeted and plate-matched control wells using a two-sided Mann–Whitney U test. Wells with non-finite feature values were excluded before analysis.

For each comparison, the numbers of target and control wells, median morphology values, median differences, Mann–Whitney U statistics, and corresponding p values were recorded, and boxplots with individual well-level observations were generated for visualization.

#### Expression–morphology associations

Genes with significant cis-eQTL support in the donor cohort were tested for association between measured expression and morphology using linear regression.

Both expression and morphology traits were mean-centered before analysis. False discovery rate was controlled separately for each morphology feature.

#### Variant–gene–morphology associations

Genes showing significant expression–morphology associations were linked to their strongest cis-eQTL variants using previously generated cis-eQTL summary statistics.

Reported associations include variant effect sizes, standard errors, and multiple-testing-adjusted significance together with corresponding expression–morphology statistics.

These statistical chains were interpreted as variant–gene–morphology associations and were not considered evidence of causal mediation.

### QUANTIFICATION AND STATISTICAL ANALYSIS

Prediction accuracy was quantified using the coefficient of determination (R²).

Feature-level significance in donor analyses was evaluated using empirical permutation testing (5,000 permutations). One-sided empirical p values were calculated using +1 smoothing and adjusted using the Benjamini–Hochberg procedure.

Shuffled-pair analyses used 1,000 random permutations.

Gene-expression–morphology associations were evaluated using ordinary least-squares linear regression.

Pearson correlation coefficients were used to quantify marginal gene–morphology relationships. CRISPR perturbation analyses were evaluated using the Mann–Whitney U test.

Unless otherwise stated, statistical significance was defined as FDR-adjusted q<0.05.

Analyses were performed using Python 3.11 with NumPy (v1.26), pandas (v2.2), SciPy (v1.11), scikit-learn (v1.5), PyTorch (v2.3), and Matplotlib (v3.9).

Exact sample sizes (n), statistical tests, and definitions of biological units are provided in the corresponding figure legends.

## ADDITIONAL RESOURCES

None.

